# p53 enables phospholipid headgroup scavenging

**DOI:** 10.1101/2024.06.07.597917

**Authors:** Jossie J. Yashinskie, Xianbing Zhu, Grace McGregor, Katrina Paras, Benjamin T. Jackson, Abigail Xie, Richard Koche, Christian Metallo, Lydia W.S. Finley

## Abstract

Changes in cell state are often accompanied by altered metabolic demands, and homeostasis depends on cells adapting to their changing needs. One major cell state change is senescence, which is associated with dramatic changes in cell metabolism, including increases in lipid metabolism, but how cells accommodate such alterations is poorly understood. Here, we show that the transcription factor p53 enables recycling of the lipid headgroups required to meet the increased demand for membrane phospholipids during senescence. p53 activation increases supply of phosphoethanolamine (PEtn), an intermediate in the Kennedy pathway for *de novo* synthesis of phosphatidylethanolamine (PE), by transactivating genes involved in autophagy and lysosomal catabolism that enable membrane turnover. Disruption of PEtn conversion to PE is well-tolerated in the absence of p53 but results in dramatic organelle remodeling and perturbs growth and gene expression following p53 activation. Consistently, CRISPR-Cas9-based genetic screens reveal that p53-activated cells preferentially depend on genes involved in lipid metabolism. Together, these results reveal lipid headgroup recycling to be a homeostatic function of p53 that confers a cell-state specific metabolic vulnerability.

## Introduction

To maintain homeostasis, cells must balance nutrient supply and demand. As cells transit through different cell states, their demand for metabolic products including lipids, nucleotides, proteins and energy may change, necessitating adaptations in nutrient supply or reorganization of metabolic networks to meet demand^1, 2^. For example, rapidly proliferating cells, like cancer cells, must continually duplicate biomass, leading to the ubiquitous model that proliferating cells will necessarily increase uptake and metabolism of nutrients to support enhanced biosynthetic demands^3–5^. While this model is a useful approximation for the metabolism of proliferative cells, it misses important nuances. For example, ATP production is rarely limiting for cell proliferation; rather, excessive ATP production may inhibit cell proliferation^6–8^. Likewise, lipid demand does not necessarily increase with proliferation. Cells synthesize new membranes at equivalent rates, regardless of proliferation; the only difference is whether or not the new membrane is used to increase total membrane pools (as in dividing cells) or replace old membranes (as in non-dividing cells)^9^. Understanding the particular metabolic requirements of distinct cell states is a prerequisite for designing effective strategies to target metabolism to eliminate malignant cell types.

One way that cells adapt to changing nutrient supply or demand is through transcription factors that coordinate gene expression programs to maintain metabolic homeostasis. During normal differentiation, lineage-defining transcription factors induce cell state changes that are often accompanied by metabolic rewiring to support changing metabolic demands^1^.

Transcription factors also respond to acute stresses that signal metabolic imbalances. For example, diverse nutrient stresses activate the tumor suppressor p53, which in turn alters expression of an array of metabolic genes that facilitate cellular adaptation and survival^10, 11^. By inducing transcriptional programs that favor nutrient oxidation over rampant glycolysis, p53 increases the efficiency by which cells convert available nutrients into ATP, thus maintaining continuous energy production despite restricted nutrient supply^10^. Under conditions of amino acid restriction, p53 balances nutrient supply and demand by increasing *de novo* amino acid synthesis while inhibiting pathways that consume amino acids^12, 13^. Notably, p53 activation also triggers changes in cell state: depending on the context, p53 activation is associated with cell cycle arrest, senescence, and cell death^11, 14^. Each of these diverse outputs will likely change cellular metabolic demands, but which metabolic resources become limiting for these altered demands, and whether p53 helps cells adapt to these cell-state dependent limitations, is poorly understood. Here, we leveraged unique cellular models of p53 activation to determine how p53 balances nutrient supply and demand during cellular senescence.

## Results

### Cells accumulate phosphoethanolamine following p53 stabilization

Because p53 is a powerful tumor suppressor, many cultured cell lines have mutated or repressed p53 or its downstream effectors, complicating efforts to study the effects of p53 on metabolism in cell culture. To circumvent this limitation, we used cells derived from a mouse model of pancreatic ductal adenocarcinoma (PDAC) in which p53 was inactivated by a doxycycline (dox)-inducible shRNA targeting *Trp53.* Here, pancreas-specific expression of oncogenic Kras (LSL-Kras^G12D^) and an mKate2-linked rtTA tet-on allele combined with dox-induced suppression of p53 allowed for formation of “KP^sh^” PDAC tumors from which we generated cell lines^15^. In KP^sh^ cells, dox withdrawal results in rapid p53 stabilization that is associated with robust induction of p53-dependent gene expression programs, growth arrest, senescence, and tumor suppression, demonstrating that these cells retain an intact p53 response^15^ (Fig. 1a). To gain a comprehensive view of metabolic changes associated with p53 activation, we performed liquid chromatography-mass spectrometry in KP^sh^ cells grown with or without dox for 6 days. Consistent with previous reports, p53 activation triggered an increase in alpha-ketoglutarate (αKG)^15^ and a decrease in many metabolites associated with nucleotide metabolism, a feature of oncogene-induced senescence^16^ (Fig. 1b and Extended Data Fig. 1a).

**Figure 1.**
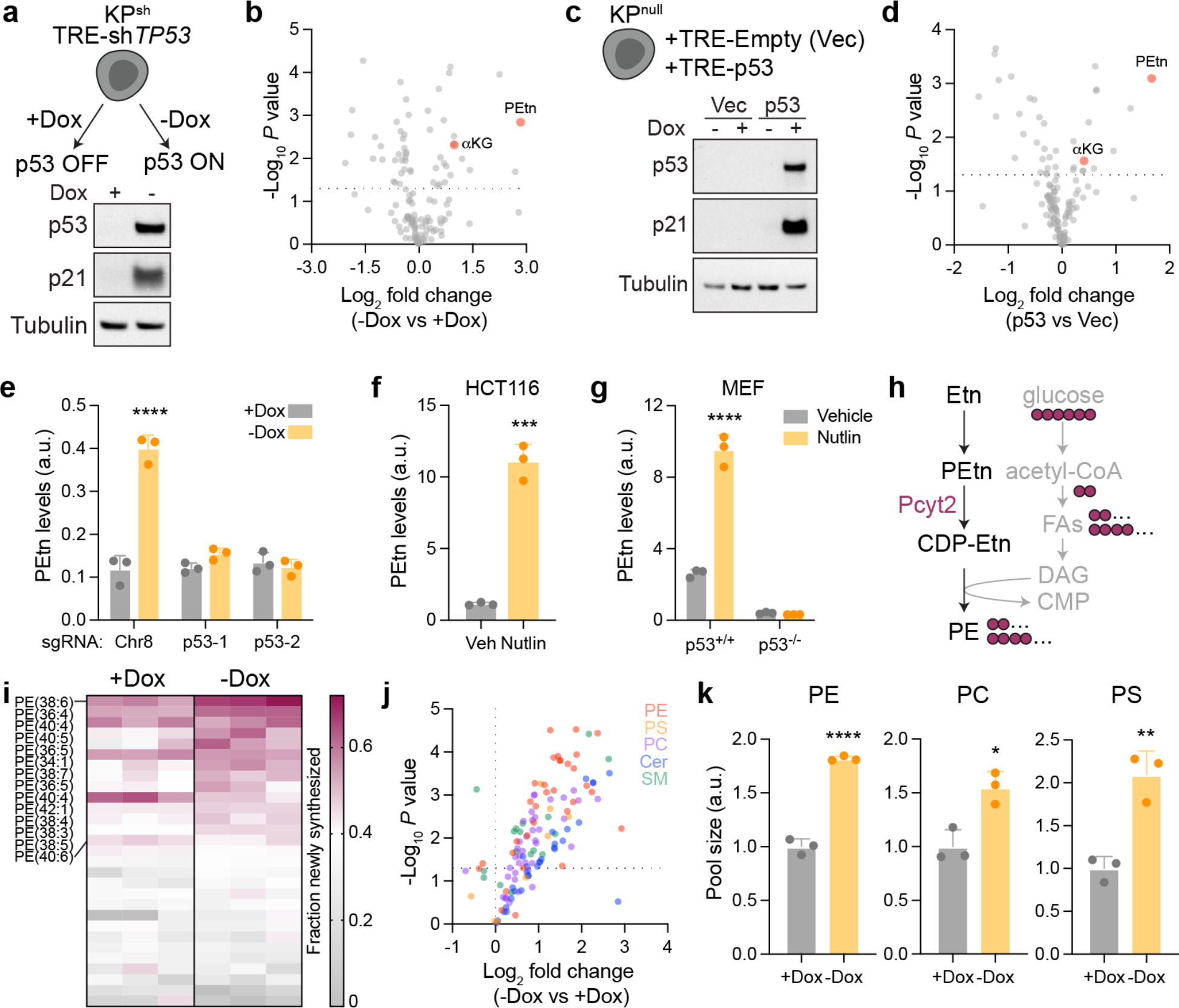
p53 activation increases phosphoethanolamine and membrane phospholipids. **a,** Schematic (top) and western blot (bottom) of KP^sh^ cells cultured with or without doxycycline (dox) for 6 days. **b,** Volcano plot showing log2 fold change in metabolite abundance in KP^sh^ cells with endogenous p53 activation (- Dox) compared to cells with silenced p53 (+Dox). Phosphoethanolamine (PEtn) and alpha-ketoglutarate (αKG) are highlighted. **c,** Schematic (top) and western blot (bottom) of KP^flox^RIK cells expressing dox-inducible vectors cultured with or without dox for 2 days. **d,** Volcano plot showing log2 fold change in metabolite abundance in KP^flox^RIK cells described in (**c**). **e,** PEtn levels in KP^sh^ cells edited with the indicated sgRNA and cultured with or without dox for 6 days. **f,g,** Steady-state levels of PEtn in HCT116 cells (**f**) or wild-type p53 (p53^+/+^) and p53 null (p53^-/-^) MEFs (**g**) cultured with DMSO (vehicle) or 5 μM Nutlin-3a (Nutlin) for 48 h. **h,** Schematic depicting *de novo* synthesis of phosphatidylethanolamine (PE) via the Kennedy pathway. [U-^13^C]glucose provides carbons (purple) that contribute to fatty acids (FAs) used in *de novo* lipid synthesis. Etn, ethanolamine; DAG, diacylglycerol. **i,** Heatmap depicting fraction of newly synthesized PE species in KP^sh^ cells cultured with or without dox for 6 days and [U-^13^C]glucose for 24 h. **j,** Volcano plot showing log2 fold change in phospholipid abundance following endogenous p53 activation in KP^sh^ cells cultured with or without dox for 6 days. Lipid species are color coded by lipid class. PS, phosphatidylserine; PC, phosphatidylcholine; Cer, ceramide; SM, sphingomyelin. **k,** Relative pool size of major lipid classes in KP^sh^ cells cultured on or off dox for 6 days. All experiments were repeated twice with similar results. Statistical significance was assessed by unpaired, two-tailed Student’s *t-*test (**b, d, f, k**) or two-way ANOVA with Sidak’s multiple comparisons post-test (**e, g**). Data are mean ± SD, *n* = 3 independent replicates. Dotted line indicates significance threshold of *P <* 0.05.

For an orthogonal cell system, we engineered rtTA-expressing cells derived from Kras^G12D^, p53 null mouse PDAC tumors (designated KP^flox^RIK) to express dox-inducible constructs containing either wild-type p53 (p53^WT^) or an empty vector. As we previously reported^15^, this system allows for rapid dox-inducible activation of p53-dependent gene expression (Fig. 1c). Despite the differences between the two systems (in one, p53 is induced by dox withdrawal; in the other, p53 is induced by dox addition), the metabolic changes were highly concordant, with αKG induced and nucleotides depleted following p53 activation in KP^flox^RIK cells (Fig. 1d and Extended Data Fig. 1b).

Across both systems, phosphoethanolamine (PEtn) emerged as the most highly induced metabolite following p53 activation (Fig. 1b, d). CRISPR/Cas9-mediated disruption of *Trp53* in KP^sh^ cells eliminated the ability of dox to induce PEtn, further confirming PEtn as a specific output downstream of p53 activation in KP^sh^ cells (Fig. 1e, Extended Data Fig. 1c). This effect was not limited to PDAC cells, as Nutlin-3a, which stabilizes p53 by inhibiting the p53-MDM2 interaction^17^, also induced PEtn in the colorectal cancer line HCT116 (Fig. 1f, Extended Data Fig. 1d). Similar results were observed in mouse embryonic fibroblasts (MEFs) expressing wild-type p53, but not in p53-null MEFs (Fig. 1g, Extended Data Fig. 1e). As HCT116 cells harbor a Kras mutation, while MEF cells do not, these results demonstrate that p53 induces PEtn accumulation independent of lineage or oncogenic mutation.

PEtn is an intermediate in the CDP-ethanolamine (Etn) arm of the Kennedy pathway for *de novo* phospholipid synthesis (Fig. 1h). An accumulation of a pathway intermediate can reflect increased influx to a pathway or downstream pathway stalling. To distinguish between these two possibilities, we cultured KP^sh^ cells in the presence of [U-^13^C]glucose and quantified isotope enrichment in phosphatidylethanolamine (PE), which lies downstream of PEtn in the CDP-Etn pathway (Fig. 1h). For most PE species, the fraction that was newly synthesized from [U- ^13^C]glucose increased following p53 activation, indicating that p53 did not impair and might even increase *de novo* phospholipid biosynthesis (Fig. 1i). In support of this conclusion, p53 activation increased the abundance of almost every membrane-associated lipid, including phosphatidylcholine (PC), phosphatidylserine (PS), sphingomyelin (SM) and ceramide (Cer) (Fig. 1j). Indeed, the total pools of major membrane phospholipids—PE, PC, PS—were significantly elevated following p53 activation (Fig. 1k). Together, these data show that cells increase synthesis and abundance of phospholipids following p53 activation.

### p53 increases ethanolamine supply

The increase in PE pools and synthesis, alongside elevated PEtn, indicate that the CDP-Etn arm of the Kennedy pathway is induced following p53 activation. *De novo* PE synthesis requires both newly synthesized acyl chains as well as ample supply of ethanolamine, which forms the lipid headgroup of PE. Neither ethanolamine nor choline (Cho), which forms the headgroup for PC and initiates the CDP-Cho arm of the Kennedy pathway, have direct synthesis pathways in mammalian cells. Rather, cells acquire ethanolamine and choline from the circulation where they are present at approximately 2 μM or 10 μM, respectively, in humans^18^. In control cells, PEtn levels are directly proportional to extracellular ethanolamine levels, rising linearly across a range of ethanolamine concentrations that spans to 10-fold above physiological levels (Fig. 2a). These results support the finding that ethanolamine uptake through its transporter FLVCR1 is likely driven by the concentration gradient between the cytosol and the extracellular space^19^ and indicate that intracellular PEtn levels are constrained by ethanolamine supply.

**Figure 2.**
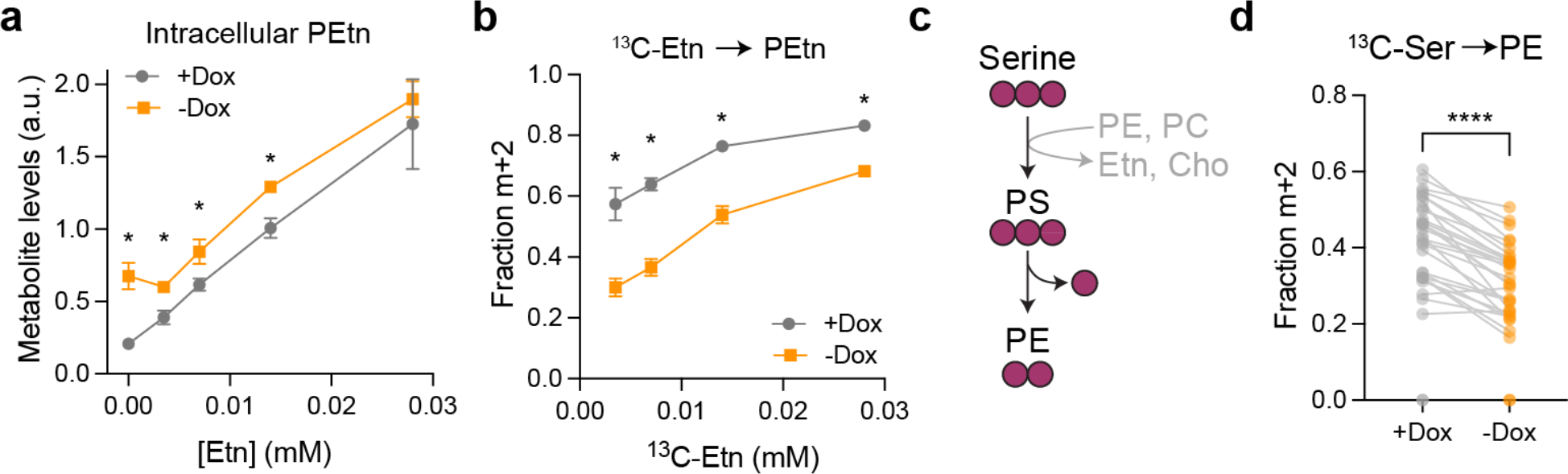
Phosphoethanolamine is limited by exogenous ethanolamine in p53-silenced cells. a, Intracellular phosphoethanolamine (PEtn) in KP^sh^ cells cultured with or without dox for 6 days with indicated concentrations of ethanolamine (Etn). **b,** Fraction of intracellular PEtn derived from exogenous [1,2-^13^C]Etn in KP^sh^ cells grown on or off dox for 6 days. **c,** Simplified schematic of [U-^13^C]serine labeling to phosphatidylethanolamine (PE). **f.** Dot plot of fractional m+2 enrichment of individual PE species in KP^sh^ cells cultured with or without dox for 6 days and [U-^13^C]serine for 24 h. Data are presented as mean (**d**) or mean ± S.D. (**a, b**) of *n* = 3 independently treated wells. Statistical significance was assessed by multiple unpaired *t*-tests with discovery determined using the two-stage step-up procedure of Benjamini, Krieger, and Yekutieli with asterisk (*) indicating false discovery rate < 1%; (**a, b**), or two-tailed paired *t-*test (**f**). Data are mean ± SD, *n* = 3 independent replicates.

Similarly, cells with active p53 also increased PEtn in response to extracellular ethanolamine availability with one major distinction: p53-activated cells were able to maintain PEtn pools even in the complete absence of exogenous ethanolamine (Fig. 2a). These results indicate that p53-activated cells have an additional source of PEtn beyond exogenous ethanolamine. To further test this possibility, we incubated cells with [1,2-^13^C]Etn, which generates m+2 labelled PEtn. Across a wide range of concentrations, [1,2-^13^C]Etn robustly labeled intracellular PEtn in p53-silenced cells (Fig. 2b). In contrast, PEtn was less labeled by [1,2-^13^C]Etn in p53 activated cells at all concentrations, confirming exogenous ethanolamine is not the only source of PEtn in these cells (Fig. 2b). Indeed, robust PEtn labeling of >40% in p53- activated cells was only achieved by supraphysiological levels of [1,2-^13^C]Etn (Fig. 2b). These results demonstrate that increased uptake and incorporation of exogenous ethanolamine cannot explain the elevated PEtn levels in p53-activated cells.

We next asked whether p53-activated cells used an alternative to the Kennedy pathway to synthesize PE. Cells can also generate PE by decarboxylating PS in mitochondrial membranes through the action of phosphatidylserine decarboxylase (PISD)^20, 21^. Activity of this pathway can be monitored by incubating cells with [U-^13^C]serine, which is incorporated into PS via base exchange and then decarboxylated to form m+2 labelled PE (Fig. 2c). Notably, despite increased PE synthesis in p53-activated cells, serine labeling to the PE headgroup was lower regardless of the PE species (Fig. 2d). Thus, p53-activated cells increase PE synthesis through routes other than PS decarboxylation. Taken together, these results demonstrate that p53 enables cells to acquire the ethanolamine lipid headgroup from sources beyond exogenous ethanolamine or serine.

### PEtn accumulation is partially dependent on p53-mediated autophagy

The above results indicate that p53 induces a cell-autonomous route of PEtn production for PE synthesis. To uncover the pathway(s) enabling PEtn production, we engineered KP^flox^RIK cells to express dox-inducible forms of p53 with mutations in one or both transactivation domains (TAD) to determine whether PEtn accumulation required the ability of p53 to transcriptionally activate downstream target genes. As expected based on previous work^22^, p53- dependent gene induction was largely preserved in TAD2 mutants, blunted in TAD1 mutants, and eliminated in TAD1/2 double mutants (Extended Data Fig. 3a, b). PEtn induction followed a similar pattern, with TAD2 mutants fully able to induce PEtn following transgene activation while TAD1 and TAD1/2 mutants were unable to accumulate PEtn despite robust p53 expression (Fig. 3a). Thus, PEtn accumulation requires p53-dependent gene transactivation.

**Figure 3.**
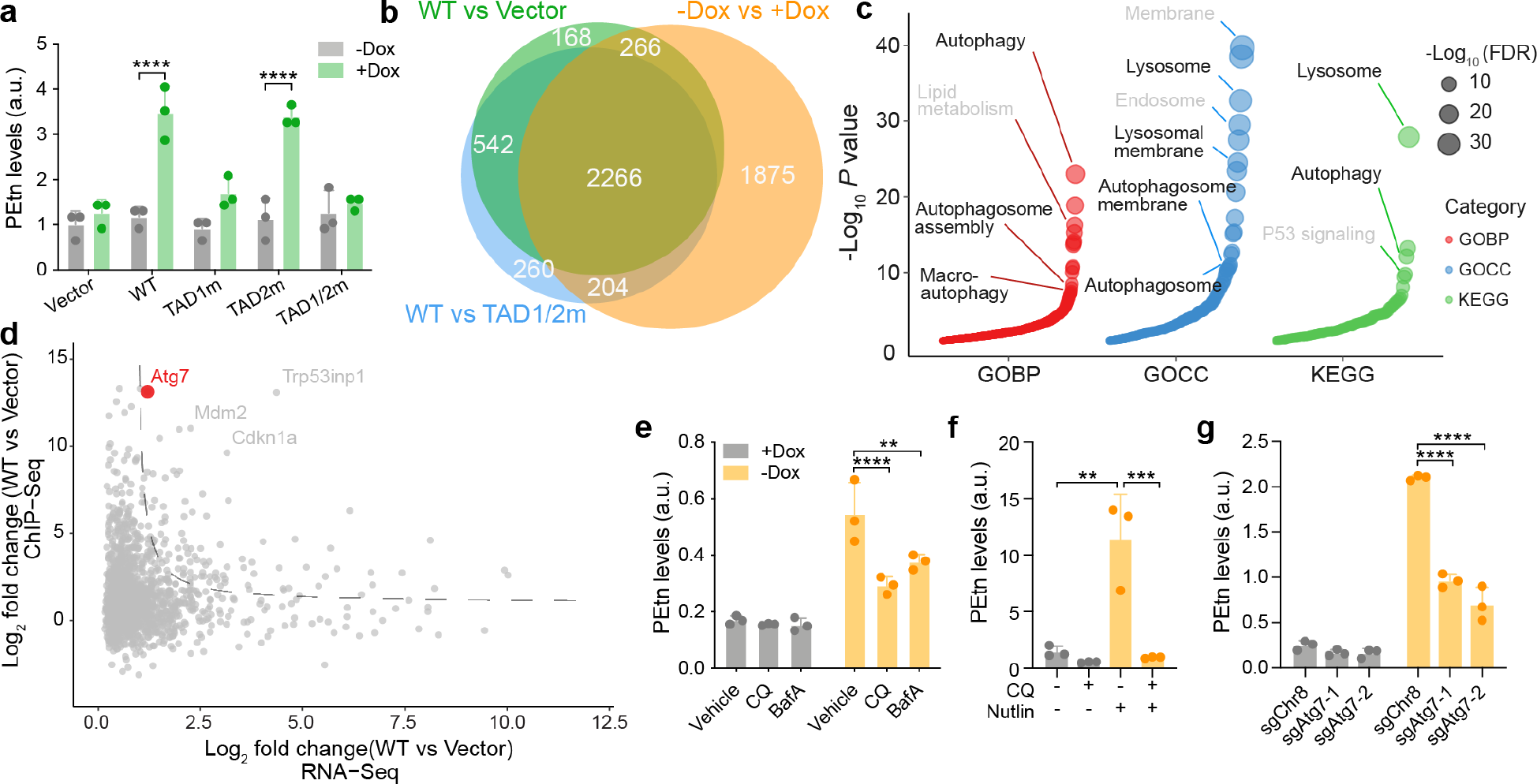
Transactivation of autophagy genes facilitates ethanolamine release by p53. a, Phosphoethanolamine (PEtn) levels in KP^flox^RIK cells engineered to express doxycycline (dox)-inducible vectors containing wild-type p53 (WT) or p53 with mutations (m) in the first or second transactivation domains (TAD) and grown with or without dox for 2 days. **b,** Venn diagram displaying the overlap of genes significantly activated (log2 fold change > 0; *P* < 0.05) in three sets of comparative analyses: WT vs Vector KP^flox^RIK cells, WT vs. TAD1/2m KP^flox^RIK cells, and -Dox vs. +Dox KP^sh^ cells. **c,** Bubble plot of pathway overrepresentation for genes induced in all 3 conditions depicted in Venn diagram in (**b**). GOBP, Gene Ontology Biological Process; GOCC, Gene Ontology Cellular Component; KEGG, Kyoto Encyclopedia of Genes and Genomes. **d,** Scatter plot comparing changes in gene expression with changes in p53 binding in KP^flox^RIK cells expressing wild-type p53 vs. vector control. Dotted line indicates threshold of log2 fold change > 1, with a curvature setting of 1.5. **e,** PEtn levels in KP^sh^ cells cultured with or without dox for 6 days and DMSO (Vehicle), 20 μM chloroquine (CQ), or 50 nM bafilomycin A1 (BafA) for the final 24 h. **f,** PEtn levels in HCT116 cells treated with DMSO (Vehicle) or 5 μM Nutlin-3a (Nutlin) for 48 h with or without 20 μM chloroquine (CQ) for the final 24 h. **g,** PEtn levels in control and *Atg7*-edited KP^sh^ cells cultured with or without dox for 6 days. Data are presented as mean ± S.D. of *n* = 3 independently treated wells with individual data points shown (**a**, **e**-**g**). Statistical significance was assessed by two-way ANOVA with Sidak’s multiple comparisons post-test (**a**), 2-way ANOVA with Dunnett’s multiple comparisons post-test (**e, g**), or 1-way ANOVA with Tukey’s multiple comparisons post-test (**f**).

To establish the common transcriptional signature associated with PEtn accumulation, we identified genes induced by wild-type p53 in both KP^sh^ and KP^flox^RIK cells (Fig. 3b). Of the 2226 genes induced by wild-type p53 in comparison with p53-silenced, -mutant, or -null cells, genes related to lysosomes and autophagy were highly overrepresented (Fig. 3c). Inspection of lysosome and autophagy gene sets revealed that the vast majority of genes in these pathways were induced by wild-type and TAD2 mutant, but not TAD1 mutant or TAD1/2 double mutant p53 (Extended Data Fig. 3c). To distinguish genes that are direct targets of p53, we profiled genomic localization of p53 using chromatin immunoprecipitation followed by sequencing (ChIP-seq). Integrating ChIP-seq and RNA-seq datasets highlighted genes both bound and activated by p53 (Fig. 3d). These genes include well-described p53 targets (e.x., *Cdkn1a*) as well as *Atg7*, which encodes an E1-like enzyme that commences the multistep process of lipidation in autophagy and has previously been reported to be a direct target of p53 (ref^23, 24^) (Extended Data Fig. 3d).

Autophagy is a degradative process that targets intracellular components to the lysosome to maintain cellular homeostasis^25^. The process of autophagy also turns over cellular membranes, leading us to hypothesize that increased autophagic flux in p53-activated cells provides an additional avenue for cells to acquire PEtn. Consistent with previous reports that p53 activates autophagy^23, 26^, p53 induction increased LC3-II production, and this effect was exacerbated by addition of bafilomycin A1 (BafA1), which inhibits lysosomal acidification and blocks autophagic turnover of LC3-II, confirming increased autophagic flux in p53-activated cells (Extended Data Fig. 3e). To determine whether lysosomal catabolism provides a source of PEtn, we incubated KP^sh^ cells with and without inhibitors of lysosomal acidification, BafA1 and chloroquine (CQ). Neither inhibitor reproducibly affected PEtn levels in control cells; in contrast, both inhibitors blunted the increase in PEtn following p53 activation (Fig. 3e).

Similarly, the increase in PEtn induced by Nutlin-3a in HCT116 cells was reversed by CQ (Fig. 3f). Next, we generated clonal KP^sh^ cell lines with genetic disruption of *Atg7* to test the hypothesis that ATG7 links p53 activation to phospholipid headgroup recycling (Extended Data Fig. 3f). Like CQ and BafA1, *Atg7* disruption blunted the p53-mediated increase in PEtn levels but had no effect in p53-silenced cells (Fig. 3g). Together, these results demonstrate that induction of genetic networks involved in lysosomes/autophagy, including transactivation of *Atg7*, facilitate PEtn accumulation in p53-activated cells.

### Ethanolamine recapture supports fitness in p53-activated cells

We next examined the impact of PEtn accumulation to p53-activated cells. PEtn is a substrate for phosphoethanolamine cytidylyltransferase (Pcyt2), which catalyzes the formation of CDP-ethanolamine from CTP and PEtn in the Kennedy pathway (Fig. 1h). Genetic disruption of *Pcyt2* increased PEtn even in control, p53-silenced cells, while further increasing PEtn in cells with active p53, consistent with elevated flux through the CDP-Etn arm of the Kennedy pathway in cells with active p53 (Fig. 4a). Re-expression of *Pcyt2* cDNA in *Pcyt2*-deficient cells restored PEtn levels to baseline, providing a genetic system to uncouple PEtn accumulation from flux through the Kennedy pathway for *de novo* PE synthesis (Fig. 4a). In control cells, *Pcyt2* disruption had a modest impact on cell proliferation and gene expression programs, demonstrating that PEtn accumulation beyond levels present in p53-activated cells is not sufficient to impair proliferation or induce significant transcriptional responses (Fig. 4b, Extended Data Fig. 4a). In contrast, p53-activated cells displayed a distinct transcriptional response to PCYT2 disruption and further slowed proliferation in the absence of PCYT2 (Fig. 4b, Extended Data Fig. 4a). Collectively, these results demonstrate that p53 activation is associated with a cell-state specific dependency on PCYT2 in the Kennedy pathway.

**Figure 4.**
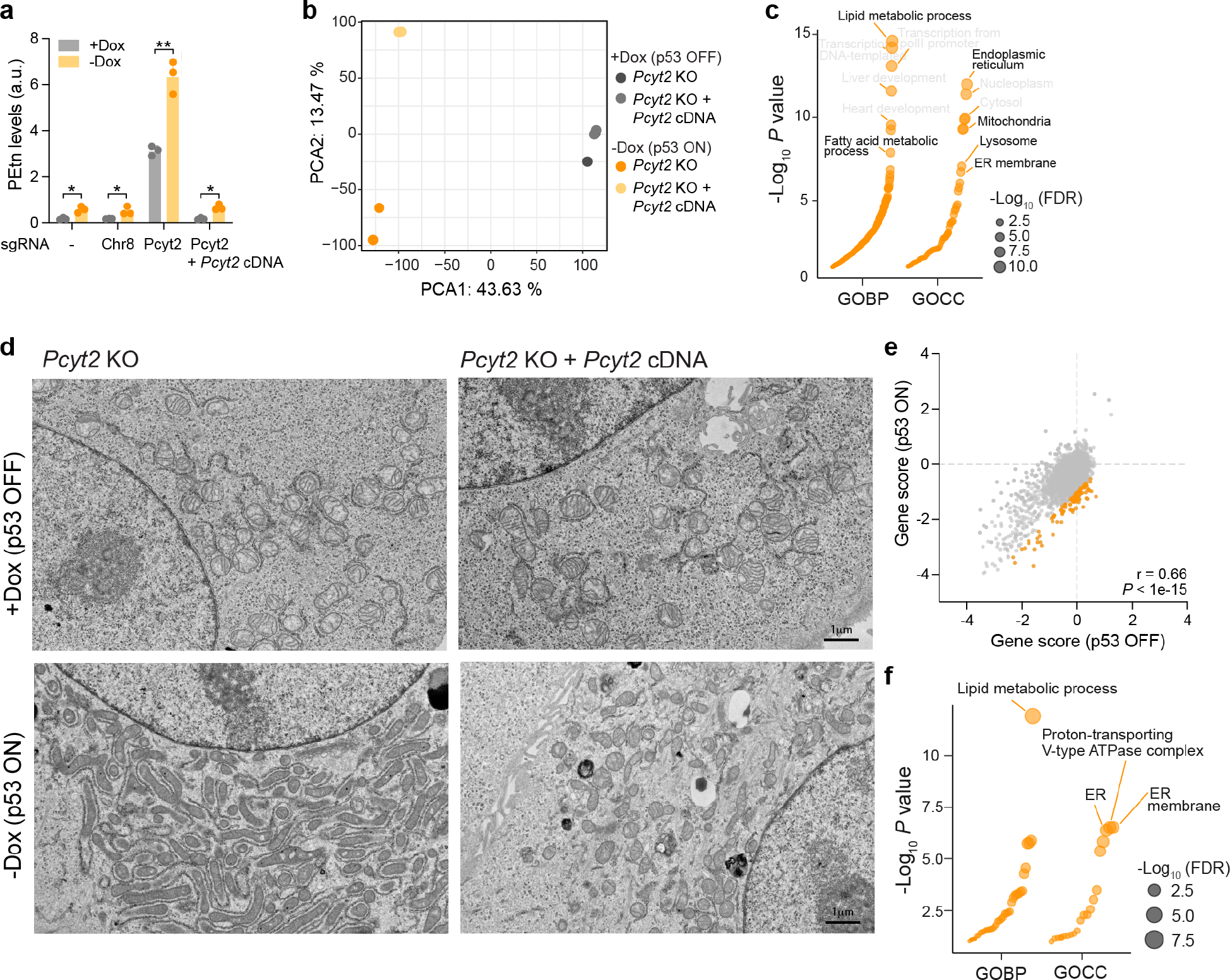
p53 activation increases reliance on genes involved in lipid metabolism. a, Phosphoethanolamine (PEtn) levels in parental KP^sh^ and clonal KP^sh^ cells expressing sgRNA and complemented with guide-resistant *Pcyt2* cDNA as indicated cultured on or off dox for 6 days. **b,c,** Principal component analysis (PCA) of RNA-Seq (**b**) and bubble plot of pathway overrepresentation for genes that are upregulated in KP^sh^ *Pcyt2*- knockout cells relative to cells complemented with sgRNA-resistant *Pcyt2* cDNA following 6 days dox withdrawal to activate p53 (**c**). **d,** Representative transmission electron microscope images from *Pcyt2*-knockout KP^sh^ cells complemented with guide-resistant *Pcyt2* cDNA cultured on or off dox for 6 days. **e,** Scatter plot comparing CRISPR gene scores KP^sh^ cells cultured with or without dox for 10 days. Genes more essential following p53 activation ([gene score -dox] – [gene score +dox] < -1) are highlighted in orange. Pearson correlation *r* = 0.66, *P* < 1 x 10^-15^. All data available in Supplementary Table 2. **f,** Bubble plot of pathway enrichment for differentially essential genes highlighted in orange in (**e**). GOBP, Gene Ontology Biological Process; GOCC, Gene Ontology Cellular Component. Data are presented as mean ± S.D. of *n* = 3 independently treated wells with individual data points shown and statistical significance was assessed by Holm-Šídák method for multiple unpaired *t-*tests. Adjusted *P* values: * < 0.05, ** < 0.01 (**a**).

To dissect the consequences of PCYT2 deficiency in p53-activated cells, we analyzed all genes that were differentially expressed in *Pcyt2* KO vs *Pcyt2* KO + *Pcyt2* cDNA-expressing cells (Extended Data Fig. 4b). Pathway enrichment analysis of transcripts upregulated in PCYT2-deficient cells revealed enrichment for genes encoding components of lipid metabolism as well as endoplasmic reticulum (ER), mitochondria, and lysosomes (Fig. 4c). Motif analysis of upregulated transcripts identified several transcription factors whose targets were overrepresented among genes induced following PCYT2 depletion, including transcription factors involved in lysosomal biogenesis (MITF, TFE3) and lipid or fatty acid metabolism (HNF1α, HNF1β, PPARα, LXRα) (Extended Data Fig. 4c). These results suggest coordinated transcriptional responses to PCYT2 deficiency that is specific to cells with activated p53.

To gain insight into how cells adjust to Kennedy pathway disruption, we performed electron microscopy to visualize PCYT2-deficient cells with or without p53 activation.

Consistent with the modest changes in growth and gene expression, *Pcyt2*-deficient cells exhibited little noticeable alteration compared to those expressing *Pcyt2* cDNA so long as p53 remained silenced (Fig. 4d). Likewise, p53 activation itself induced few dramatic changes in PCYT2-expressing cells, although scattered lipid droplets became apparent in line with previous work demonstrating that p53 activation and senescence are associated with induction of lipid droplets in some contexts^27, 28^ (Fig. 4d). In contrast, p53 activation in PCYT2-deficient cells induced dramatic cellular remodeling, with mitochondria becoming surrounded by curved sheets of rough ER (Fig. 4d). Similar patterns of mitochondria enwrapped by ER are induced during fasting in several mouse tissues and proposed to facilitate lipid exchange to sustain fasting lipid homeostasis^29^. Notably, the ER is the site of PE synthesis downstream of PCYT2 in the Kennedy pathway^30^. Given that mitochondria—and in particular, mitochondrial-ER interactions—are sites of PE synthesis from PS decarboxylation^29, 31^, our results are consistent with a model in which ER approach mitochondria to acquire PE when the cellular demand for PE cannot be met by the Kennedy pathway alone.

Together, these results demonstrate that p53 activation increases demand for new phospholipid synthesis which is met in part through *de novo* synthesis through the Kennedy pathway with autophagy and lysosomal catabolism supplying PEtn for lipid headgroups. We next aimed to determine whether metabolic networks involved in lipid synthesis and/or autophagy contributed to cell fitness in p53-activated cells. To this end, we performed CRISPR- Cas9-based genetic screens in KP^sh^ cells using a metabolism-focused sgRNA library^32^. Cells were transduced with the Cas9-sgRNA library in the presence of dox and, following selection, grown in the presence or absence of dox for 10 days. Gene scores were determined by comparing final sgRNA abundance at day 10 with initial sgRNA abundance at day 0. Overall, cells grown with and without dox exhibited highly correlated gene scores, indicating that the requirement for many metabolic genes remains largely consistent regardless of p53 activation (Fig. 4e).

Nevertheless, differences emerged, with some genes becoming relatively more or less essential following p53 activation. Notably, genes demonstrating increased essentiality following p53 activation were strongly enriched for genes encoding components of lipid metabolic processes, cellular membranes, or the V-type ATPase required for lysosomal acidification (Fig. 4f). Of note, the sgRNA library is limited to metabolic genes, and thus does not include sgRNA targeting many aspects of autophagy or lysosomal biology beyond those encoding the V-type ATPase.

We reasoned that these CRISPR-Cas9 screens provided an opportunity to determine which p53-regulated metabolic genes contribute to cell fitness following p53 activation. Genes induced by p53 showed a weak, yet statistically significant, tendency to become more essential following p53 activation (Extended Data Fig. 4d). In fact, many genes induced by p53 appeared to antagonize fitness in p53 activated cells, indicating that gene induction is not sufficient to infer necessity. Of the genes both induced by p53 and more essential for fitness following p53 activation, we noted strong overrepresentation of genes involved in lipid metabolism (Extended Data. Fig. 4e). Collectively, these results indicate that p53 activation both increases lipid synthesis and induces reliance on lipid metabolism to support cell fitness.

## Discussion

Here we show that p53 activation triggers increased synthesis and accumulation of phospholipids and this program of elevated lipid metabolism is required for fitness in the context of p53 activation. Notably, phospholipid synthesis requires provision of phospholipid headgroups, which cells usually acquire from circulation. In most cell culture conditions, the sole source of the ethanolamine lipid headgroup is serum; culturing cells in dialyzed serum thus deprives cells of exogenous ethanolamine. How, then, can cells meet demand for increased phospholipid synthesis? We find that cells can also acquire ethanolamine by recycling lipids through the autophagy/lysosomal system, and this process is induced following p53 activation. In many contexts, p53 helps cells adapt to conditions of nutrient limitation, thus facilitating survival under challenging conditions^10^. Our results reveal an additional mechanism by which p53 allows cells to cope with nutrient stress: by increasing autophagy which supplies headgroups for *de novo* lipid biosynthesis.

Increased lysosomal content and activity is a common feature of senescence and indeed represents the basis for the ubiquitous senescence marker, senescence-associated β-galactosidase (SA-βgal), which reads out elevated function of the lysosomal enzyme β-galactosidase^33^.

Consistently, PEtn upregulation is also a common feature of senescent cells^34^, suggesting a tight correlation between PEtn accumulation and lysosomal activity. Senescence is also broadly associated with altered lipid metabolism, including increases in both synthesis and oxidation of fatty acids, likely reflecting increased turnover of lipid pools in senescent cells^27^. During senescence, cells enlarge and accumulate organelles including lysosomes, mitochondria, and lipid droplets^27, 35–37^. Such organelle accumulation necessarily requires generation of phospholipid membranes, which in turn require the acyl chains, glycerol backbone, and phospholipid headgroup that comprise membrane phospholipids. Our results demonstrate that p53 helps cells meet these increased demands by facilitating the provision and recapture of the ethanolamine headgroup for PE synthesis. Given that the high-affinity ethanolamine transporter was only recently identified^19^, future work will be needed to identify conditions under which exogenous ethanolamine availability constrains *de novo* lipid synthesis in cells *in vivo*.

More broadly these results highlight how the metabolic demands of specific cell states require coordinated metabolic remodeling. In the case of p53 activation, induction of genes involved in autophagy/lysosomal biology enable phospholipid recycling to supply key intermediates required for increased phospholipid accumulation in senescent cells. The demand for increased phospholipids introduces cell-state specific metabolic vulnerabilities: namely, *de novo* phospholipid synthesis, tolerated in p53 null cells, becomes essential in contexts of p53 activation. Given that cells have redundant pathways to generate membrane phospholipids, including *de novo* synthesis, base exchange with existing phospholipids, or scavenging from extracellular sources, it is perhaps not surprising that disruption of one axis (here, PCYT2 in the CDP-Etn arm of the Kennedy pathway) is tolerated so long as the demand for membrane phospholipids remains low. Taken together, our work raises the possibility that disrupting lipid headgroup supply—whether via transporter uptake or recapture through the Kennedy pathway— may represent a selective liability in p53-activated cells, whether in tumors or during normal aging.

## Experimental Methods

### Cell culture

KP^sh^ and KP^flox^RIK cells were previously generated^15^. KP^sh^ cells were cultured in DMEM supplemented with 10% fetal bovine serum (FBS; Gemini) and 1 μg/ml doxycycline (Sigma) on collagen-coated plates (PurCol, Advanced Biomatrix, 0.1 mg/ml). Medium was refreshed every other day. KP^flox^RIK cells were cultured in complete DMEM supplemented with 10% FBS on collagen-coated plates and 1 μg/ml doxycycline was administered to induce transgene expression 48 h prior to harvest. HCT116 and 293T cells were obtained from ATCC. MEFs, a kind gift of John P. Morris IV and Scott Lowe (ref^38^), were maintained at 3% oxygen until seeded for experiments. Where indicated, cells were treated with DMSO (vehicle control, Sigma), 5 μM Nutlin-3a (Selleck Chemicals), 20 μM Chloroquine (Sigma), 50 nM Bafilomycin A (Sigma), or 3.5, 7, 14, or 28 μM Etn (Sigma).

### Genetically-edited cell lines

sgRNA sequences targeting *Atg7* (CACCGTCTCCTACTCCAATCCCGTG), *Pcyt2*, (CACCGCCATGATCCGGAACGGGCA) or a non-genic region on mouse chromosome 8 (Chr8, ACATTTCTTTCCCCACTGG) were cloned into lentiCRISPRv2 (Addgene, 52961). sgRNA sequences targeting *Trp53* (GAAGTCACAGCACATGACGG; GCAGACTTTTCGCCACAGCG) or a non-genic region on mouse chromosome 8 (Chr8) were cloned into the pUSEPB plasmid^15^ (gift from S. Lowe). Mouse wild-type *Pcyt2* cDNA (Horizon Discovery, MMM1013-202763346) was cloned into N174-MCS (Addgene, 81061) backbone digested with EcoRI and BamHI (New England Biolabs) using the Q5 High-Fidelity DNA Polymerase (New England Biolabs, M049) according to the manufacturer’s instructions. Site directed mutagenesis was performed according to the manufacturer’s instructions to introduce gRNA target-site mutations in wild-type *Pcyt2* using the Q5 Site-Directed Mutagenesis Kit (New England Biolabs, E0554). Primers were designed using NEBaseChangerv1.

Lentivirus was generated by co-transfection of viral vectors expressing sgRNA or cDNA of interest with packaging plasmids psPAX2 (Addgene, 12260) and pMD2.G (Addgene, 12259) into 293T cells. Viral-containing supernatants were cleared of cellular debris by 0.45 μM filtration and mixed with 8 μg/ml polybrene (Sigma). Target cells were exposed to viral supernatants for two 24 h periods before being washed, grown for 12 h in fresh media, then subjected to antibiotic selection until control cells were eliminated. Cells were maintained in antibiotic selection until used for experiments. To generate clonal lines, cells were sequentially diluted to a final concentration of 1 cell per 200 μl of complete DMEM, seeded onto 96-well tissue culture plates, and expanded to enable clonal growth. Loss of target gene expression was validated by immunoblotting or metabolic profiling.

### Western blotting

Protein lysates were extracted in 1X RIPA buffer (Cell Signaling Technology) and protein concentration was determined by BCA assay (Thermo Fisher Scientific). Samples were boiled for 5 min and 20 to 30 μg of protein were separated by SDS–polyacrylamide gel electrophoresis and transferred to nitrocellulose membranes (Bio-Rad). Membranes were blocked in 3% milk in Tris-buffered saline with 0.1% Tween 20 (TBST) and incubated at 4 °C with primary antibodies overnight. Membranes were then washed with TBST and incubated with horseradish peroxidase (HRP)-conjugated secondary antibodies for at least 4 h. Proteins were detected using Pierce ECL Western Blotting Substrate (Thermo Fisher Scientific, 32106) and imaged using HyBlot CL Autoradiography Film (Denville Scientific, E3018) and SRX-101A X-ray Film Processor (Konica Minolta). Antibodies were diluted as follows: anti-p53 (CM5) (1:500, Leica Biosystems, NCL-L-p53-CM5p), anti-p21 (F-5) (1:500, Santa Cruz Biotechnology, sc-6246), anti-PCYT2 (1:1000, Proteintech, 14827-AP), anti-LC3a/b (D3U4C) (1:1000, Cell Signaling Technology, 12741T), anti-ATG7 (1:1000, Proteintech, 10088-2-AP) and anti-Vinculin (1:10,000, Sigma, V9131).

### RNA-Seq

Total RNA was isolated from duplicate wells of 6-well plates using TRIzol (Invitrogen) according to the manufacturer’s instructions and quantified using a BioAnalyzer (Agilent). Preparation of the RNA-Seq libraries and sequencing was performed by the Memorial Sloan Kettering Cancer Center Integrated Genomics Operation. RNA-Seq files were mapped to the reference mouse genome sequence (mm10) with STAR (2.6.1c)^39^. The number of raw counts was calculated by Homer^40^ with Gencode gene annotation GTF file and normalized by Fragments Per Kilobase per Million mapped fragments (RPKM). Significant differentially expressed genes between conditions were identified using DESeq2 (version 1.19.38)^41^ with a p-value < 0.05. Motif analysis was conducted using Homer, searching from -400 to +100 relative to the transcription start site (TSS). The pathway enrichment analysis was performed using DAVID (https://david.ncifcrf.gov/home.jsp) and the enriched pathways were visualized in R (V4.3.1). The principal component analysis (PCA) and heatmap were generated using FactoMineR() and pheatmap() R package, respectively. Data used to generate heatmaps is provided in Supplementary Table 1.

### ChIP-Seq

Chromatin immunoprecipitation experiments were carried out as described^42^ with modest modifications. Cells were crosslinked with 1% formaldehyde for 10 min at room temperature. Crosslinking was quenched by adding glycine to a final concentration of 125 mM for 5 min. Cells were washed twice with cold PBS, then flash frozen and stored at -80°C for up to 1 week. Cells were processed for sonication by sequential resuspension with the following buffers: cold Lysis Buffer 1 (50 mM Hepes-KOH pH 7.5, 140 mM NaCl, 1 mM EDTA pH 8.0, 10% glycerol, 0.5% NP-40, 0.25% Triton X-100, 1× protease inhibitors (Thermo Fisher Scientific, 78438)) for 10 min at 4 °C, cold Lysis Buffer 2 (10 mM Tris-HCl pH 8.0, 200 mM NaCl, 1 mM EDTA, 0.5 mM EGTA, 1× protease inhibitors) for 10 min at room temperature, and finally, Sonication Buffer (50 mM Tris-HCl, pH 8.0, 10 mM EDTA, 0.5% SDS). Lysed cells were sonicated to generate DNA fragments with an average length of about 100-400 bp. Subsequently, 5.5 μg of p53 (1C12) antibody (Cell Signaling Technology) and 55 μL of Dynabeads Protein G (Thermo Fisher Scientific) were used for each ChIP. DNA was eluted at 65°C for 30 min with 50 mM Tris-HCl, pH 8.0, 10 mM EDTA, 1.0% SDS and incubated with 200 mM NaCl and 20 μg RNAse A at 65°C overnight for reversal of crosslinking. Protein was digested with 80 μg proteinase K at 55°C for 2 h. DNA was purified using phenol:chloroform:isoamyl alcohol (P:C:IA, Sigma). Eluted DNA was submitted to the Integrated Genomics Operation at Memorial Sloan Kettering Cancer Center for library preparation and sequencing. ChIP-Seq files were mapped to reference mouse genome sequence (mm10) using bowtie2 with default parameters.

Duplicate reads were removed using samtools^43^. Peaks in the ChIP-seq data were identified using MACS2 (ref^44^), and the reads over these peaks were quantified using Bedtools^45^. Summation of peak reads for individual genes was performed. The fold change of normalized reads between ChIP-Seq and RNA-Seq data was visualized using the ggplot2() R package, applying a log2 fold change threshold of 1 and a curvature setting of 1.5. Bigwig files of ChIP-Seq data were generated using Deeptools^46^ and ChIP-Seq tracks were visualized using Integrative Genomics Viewer^47^.

### Polar metabolite profiling and stable isotope tracing

Cells were seeded 2 days before collection in their standard culture medium in either 6-well plates (LC-MS) or 12-well plates (GC-MS). Medium was changed again at the indicated time before harvest (usually 1–24 h). Metabolites were extracted with 1 mL ice-cold 80% methanol/20% water solution supplemented with 2 μM deuterated 2-hydroxyglutarate (D-2- hydroxyglutaric-2,3,3,4,4-d5 acid, d5-2HG) as an internal standard. Immediately after, plates were transferred to -80 °C and incubated overnight. Metabolite extracts were dried in an evaporator (Genevac EZ-2 Elite).

For metabolomic profiling and stable isotope tracing analysis using gas-chromatography-mass spectrometry (GC-MS), dried extracts were resuspended in 20 μl of 40 mg/mL methoxyamine hydrochloride in pyridine, then incubated with shaking at 30°C for 1.5 h. Metabolites were further derivatized by adding 18 μl of *N*-methyl-*N*-(trimethylsilyl) trifluoroacetamide with or without 1% TMCS (Thermo Fisher Scientific) and 20 μl ethyl acetate (Sigma) and then incubated at 37°C for 30 min. Samples were analyzed using an Agilent 7890A gas chromatograph coupled to Agilent 5975C mass selective detector. The gas chromatograph was operated in splitless mode with helium gas flow at 1 mL/min. l μl of sample was injected onto an HP-5MS column and the GC oven temperature ramped from 60°C to 290°C over 25 min. Peaks representing compounds of interest were extracted and integrated using MassHunter software (Agilent Technologies) and then normalized to both the internal standard (d5-2HG) peak area and protein content of duplicate samples as determined by BCA protein assay (Thermo Fisher Scientific). Ions used for quantification of metabolite levels are as follows: d5-2HG, 354 *m*/*z* and PEtn, 188 *m*/*z*. All peaks were manually inspected and verified relative to known spectra for each metabolite. For ethanolamine isotope tracing studies: 1 h before metabolite collection, cells were washed and incubated with complete DMEM supplemented with 10% dialyzed fetal bovine serum (dFBS; Gemini), and ^12^C-ethanolamine (Sigma) or [1,2-^13^C]ethanolamine (Cambridge Isotope Labs).

Enrichment of ^13^C was determined by quantifying the abundance of the following ions: PEtn, *m/z* 188-190. Correction for natural isotope abundance was performed using IsoCor (v.2.0)^48^.

For metabolomic profiling using liquid-chromatography-mass spectrometry (LC-MS), dried extracts were resuspended in 30 μL of mobile phase A (10 mM tributylamine and 15 mM acetic acid in 97:3 water:methanol). Samples were incubated on ice for 20 min with vortexing every 5 min then cleared by centrifugation at 20,000*g* for 20 min at 4°C. Reconstituted samples were subjected to MS/MS acquisition using an Agilent 1290 Infinity LC system equipped with a quaternary pump, multisampler, and thermostatted column compartment coupled to an Agilent 6470 series triple quadrupole system with a dual Agilent Jet Stream source for sample introduction. Data were acquired in dynamic MRM mode using electrospray ionization in negative ion mode. Capillary voltage was 2000 V, nebulizer gas pressure was 45 Psi, drying gas temperature was 150°C and drying gas flow rate was 13 L/min. 5 μL of sample was injected on to an Agilent Zorbax RRHD Extend-C18 (1.8 μm, 2.1 × 150 mm) column maintained at 35°C. A 24 min chromatographic gradient was performed using mobile phase A and mobile phase B (10 mM tributylamine and 15 mM acetic acid in methanol) at a flow rate of 0.25 mL/min. Following the 24 min gradient the analytical column was backflushed for 6 min with 99% acetonitrile at a 0.8 ml/min flow rate followed by a 5 min re-equilibration step (MassHunter Metabolomics dMRM Database and Method, Agilent Technologies). Data analysis was performed with MassHunter Quantitative Analysis (v. B.09.00). Peak measurements were normalized to protein content of duplicate samples as determined by BCA protein assay (Thermo Fisher Scientific). All experiments were performed independently at least three times and a representative experiment is shown.

### Lipid metabolite profiling

For lipid extraction, cells were seeded 2 days before collection in their standard culture medium in 6-well plates. 24 h before harvest, cells were washed with PBS and incubated with glucose- and serine-free DMEM supplemented with 10% dialyzed fetal bovine serum (dFBS; Gemini), ^12^C-glucose and ^12^C-serine (Sigma) or the labeled versions of each metabolite: [U-^13^C]glucose or [U-^13^C]serine (Cambridge Isotope Labs). At harvest, cells were washed with PBS at room temperature and then lysed with 750 μl of ice-cold 5:4 methanol/water solution for 5 min on ice and scraped into Eppendorf tubes. Next, 500 μl of ice-cold chloroform (Sigma) and 50 μl of butylated hydroxytoluene (1 mg/ml in methanol; Sigma) were added to the tube, along with EquiSLASH LIPIDOMIX labelled standard (Avanti Polar Lipids). The tubes were vortexed for 5 min and centrifuged at 20,000*g* at 4°C for 5 min. The lower organic phase was collected, and 2 μl of formic acid (Sigma) was added to the remaining polar phase which was re-extracted with 500 μl of chloroform. Combined organic phases were dried, and the pellet was resuspended in 50 μl of isopropyl alcohol.

Broad spectrum lipidomics analysis was performed on a Q Exactive orbitrap mass spectrometer with a Vanquish Flex Binary UHPLC system (Thermo Fisher Scientific) using an Accucore C30, 150 x 2.1 mm, 2.6 µm particle (Thermo Fisher Scientific) column maintained at 40°C. Chromatography was performed at a flow rate of 0.2 mL/min using a gradient of 40:60 v/v water: acetonitrile with 10 mM ammonium formate and 0.1% formic acid and 10:90 v/v acetonitrile: 2-propanol with 10 mM ammonium formate and 0.1% formic acid (mobile phase B). The LC gradient ran from 30% to 43% B from 3-8 min, then from 43% to 50% B from 8-9 min, then 50-90% B from 9-18 min, then 90-99% B from 18-26 min, then held at 99% B from 26-30 min, before returning to 30% B in 6 min and held for a further 4 min. 5 μL of sample was injected.

Lipids were analyzed in positive mode using spray voltage 3.2 kV. Sweep gas flow was 1 arbitrary units, auxiliary gas flow 2 arbitrary units and sheath gas flow 40 arbitrary units, with a capillary temperature of 325°C. Full MS (scan range 200-2000 *m/z*) was used at 70,000 resolution with 1e6 automatic gain control and a maximum injection time of 100 ms. Data dependent MS2 (Top 6) mode at 17,500 resolution with automatic gain control set at 1e5 with a maximum injection time of 50 ms was used. Data was analyzed using EI-Maven software, using at least x2 fragment ions (where possible depending on lipid species) for accurate species identification, and peaks normalized to Avanti EquiSPLASH internal standard. Pool size was calculated using the normalized abundance and cell protein content. Mass isotopomer enrichment calculations were performed using a combination of in-house Matlab scripts and Escher-trace software^49^.

### CRISPR screens

The mouse metabolism library (ref^32^) was a generous gift from Kivanc Birsoy and was amplified using Endura Electro-Competent cells (Lucigen). Lentivirus was produced for the screen library by the co-transfection of the screen library plasmid with the packaging plasmids psPAX2 and pMD2.G into 293T cells as described above. 30 million KP^sh^ cells were infected at an MOI of 0.3 to maintain representation of 300X. KP^sh^ cells were maintained on doxycycline (Sigma, 1 μg/mL) until the start of the screen. Cells were selected in puromycin (Thermo Fisher Scientific, 1 μg/mL) for 7 days, after which genomic DNA samples were collected (Day 0) and cells were seeded into screen conditions. Cells were cultured in the presence or absence of doxycycline and passaged every 48 h for 10 days. Samples for genomic DNA were collected at the conclusion of the screen (Day 10). Following the screen, genomic DNA was extracted using the Blood & Cell Culture DNA Kit Midi (Qiagen) according to manufacturer instructions. PCR amplification of sgRNA inserts was performed using Ex Taq DNA Polymerase (Takara) with a total of 54 μg of input genomic DNA to maintain 300X representation. PCR products for each sample were pooled and submitted to the Integrated Genomics Operation at MSKCC for library preparation and sequencing. Reads were aligned using Rsubread (version 2.2.6)^50^, and normalized by dividing the number of reads per sgRNA by the total number of reads in each sample, multiplying by a scaling factor of 10^6^, and adding a pseudocount of 1. sgRNA scores were calculated by taking the log2 fold change of sgRNAs between conditions of interest, and gene scores were calculated by taking the median of all sgRNA scores for each gene. Data used to generate the scatterplots in the figures is provided in Supplementary Table 2.

### Electron Microscopy

Samples were processed and imaged at The Rockefeller University Electron Microscopy Resource Center. Cells were fixed in 2% glutaraldehyde in 0.08 M sodium cacodylate buffer (pH 7.2) with 2 mM CaCl2 for 4 h at room temperature followed by overnight fixation at 4°C, postfixed in 1% osmium/0.8% potassium ferricyanide in 0.1 M cacodylate buffer, followed by post-staining in 1% uranyl acetate in water, dehydration in an ethanol series, and embedding in Eponate 12 (Ted Pella, Inc). Ultrathin sections (60–65 nm) were stained with uranyl acetate and lead citrate, and images were acquired using a Tecnai G2-Spirit transmission electron microscope (FEI, Hillsboro, Oregon) operated at 120 kV, equipped with an AMT BioSprint29 digital camera (AMT, Danvers, MA).

### Statistics and reproducibility

GraphPad PRISM 10 software was used for statistical analyses unless otherwise noted. Sample size, error bars and statistical tests are reported in the figure legends. Statistical tests include unpaired two-tailed Student’s *t-*test, one-way analysis of variance (ANOVA), two-way ANOVA and Fisher’s exact test. All sequencing experiments were done once with the indicated number of biological replicates. The remaining experiments were performed independently at least two times.

### Data and reagent availability

Sequencing data supporting the findings of this study have been deposited in the Gene Expression Omnibus under the accession codes: GSE268723 (RNA-Seq) and GSE268722 (ChIP-Seq). p53 ChIP-seq in primary MEFs cells was obtained from published sources^23^ (GSE46240).

All unique reagents generated in this study will be made available upon request from the corresponding author with a completed material transfer agreement.

## Acknowledgements

We thank members of the Finley lab for discussion, Andrew Intlekofer for shared use of LC-MS instrumentation, Sara Violante and Justin Cross for assistance with LC-MS protocols, Alexander Muir for sharing protocols for measuring phosphoethanolamine, John P. Morris IV for sharing MEFs, and The Rockefeller University Electron Microscopy Resource Center for performing EM. J.J.Y. was supported by T32HD060600. K.I.P. was supported by a Bruce Charles Forbes Pre-Doctoral Fellowship (MSKCC). A.X. and B.T.J. are supported by Ruth L. Kirschstein Predoctoral fellowships (no. F30CA284711 to A.X.; no. F30HD107943 to B.T.J.), and a Medical Scientist Training Program grant from the NIGMS of the National Institutes of Health under award number T32GM007739 to the Weill Cornell/Rockefeller/Sloan Kettering Tri-Institutional MD-PhD Program. L.F. is a New York Stem Cell Foundation – Robertson Investigator and was a Searle Scholar. This work was additionally supported by grants from NYSCF, the Anna Fuller Fund, The Conquer Cancer Now Fund, the Edward Mallinckrodt Jr., Foundation, the NIH/NCI (R37 CA252305 to L.F.; R01 CA234245 to C.M.), as well as the Memorial Sloan Kettering Cancer Center Support Grant P30CA008748.

## Author contributions

J.J.Y. and L.W.S.F. conceived the study. J.J.Y. carried out experiments with assistance from X.Z, K.P., and A.X. K.P. performed the CRISPR/Cas9 screen and RNA-seq analysis with assistance from B.T.J. and X.Z. R.K. and X.Z. analyzed ChIP-Seq. G.M. performed lipidomic analysis with guidance from C.M. X.Z. assisted with data visualization. J.J.Y. and L.W.S.F. wrote the manuscript with input from all authors.

---

**Extended Data Figure 1.**
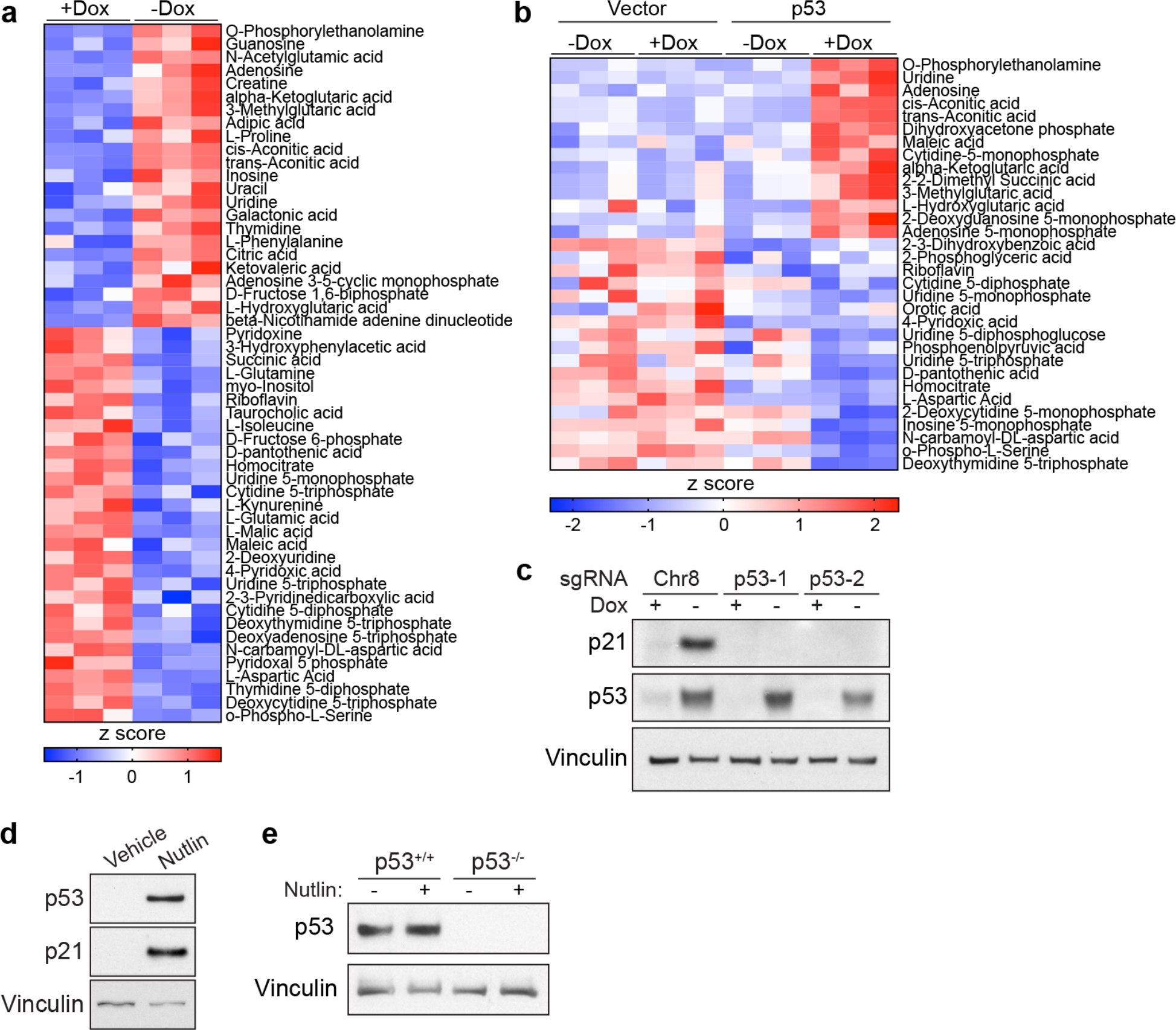
a,b, Heatmap displaying relative abundance of all metabolites exhibiting significant (*P* < 0.05) change when comparing KP^sh^ cells cultured with or without dox for 6 days (**a**) or KP^flox^RIK cells expressing dox-inducible p53 compared with vector expressing cells cultured with dox for 2 days (**b**). **c,** Western blot of KP^sh^ cells edited by the indicated sgRNA and cultured with or without dox for 6 days. **d,e,** Western blot of HCT116 cells (**d**) or wild-type p53 (p53^+/+^) and p53 null (p53^-/-^) MEFs (**e**) cultured with DMSO (vehicle) or 5 μM Nutlin-3a (Nutlin) for 48 h.

**Extended Data Figure 3.**
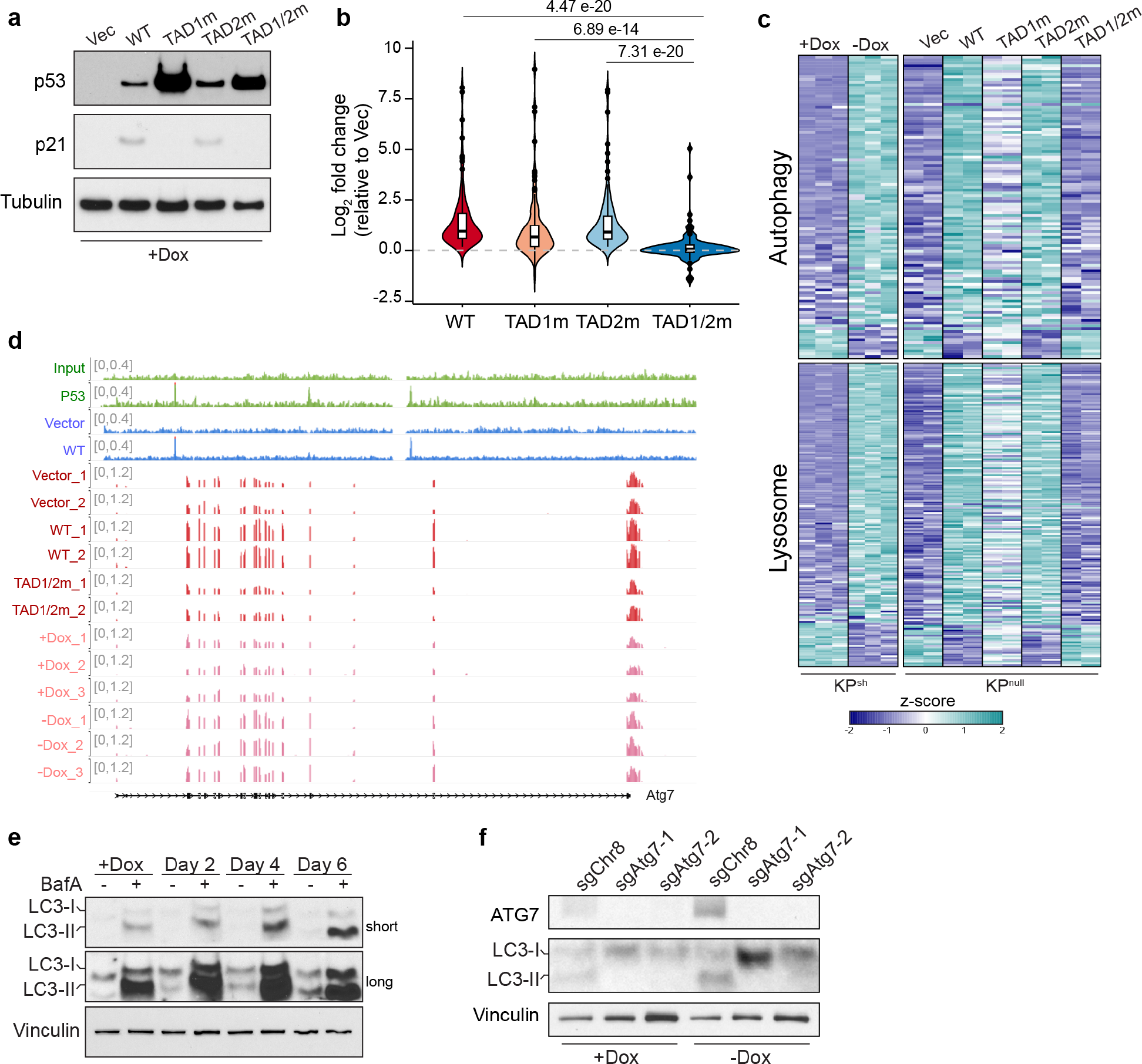
Transactivation of autophagy genes facilitates ethanolamine release by p53 a, Western blot of KP^flox^RIK cells engineered to express doxycycline (dox)-inducible vectors containing wild-type p53 or p53 with mutations (m) in the first or second transactivation domains (TAD) grown with dox for 2 days. **b,** Violin plot showing log2 fold change in expression of HALLMARK p53 pathway genes activated by wild type p53 (log2 fold change > 0; *P* < 0.05) in KP^flox^RIK cells cultured with dox for 2 days. Log2 fold change is expressed relative to cells expressing empty vector. Statistical significance was assessed by paired *t*-test using R. **c,** Heatmap showing expression of autophagy and lysosome genes in KP^sh^ cells grown on or off dox for ten days and KP^flox^RIK cells cultured on dox for two days. Genes with *P* < 0.05 in KP^sh^ cells are ranked according to fold change. **d,** Sequencing tracks for the *Atg7* locus, displaying p53 ChIP-seq (top) and RNA-seq (bottom) signals in doxorubicin-treated primary MEFs (ref^23^, green) or KP^flox^RIK cells expressing vector or wild-type p53 treated with dox for 2 days (blue). **e,** Western blot (short and long exposure) of KP^sh^ cells cultured with or without 50 nM bafilomycin A1 (BafA) for 24 h and exposed to dox withdrawal for indicated time. **f,** Western blot of clonal KP^sh^ cells expressing indicated sgRNA and cultured with or without dox for 6 days.

**Extended Data Figure 4.**
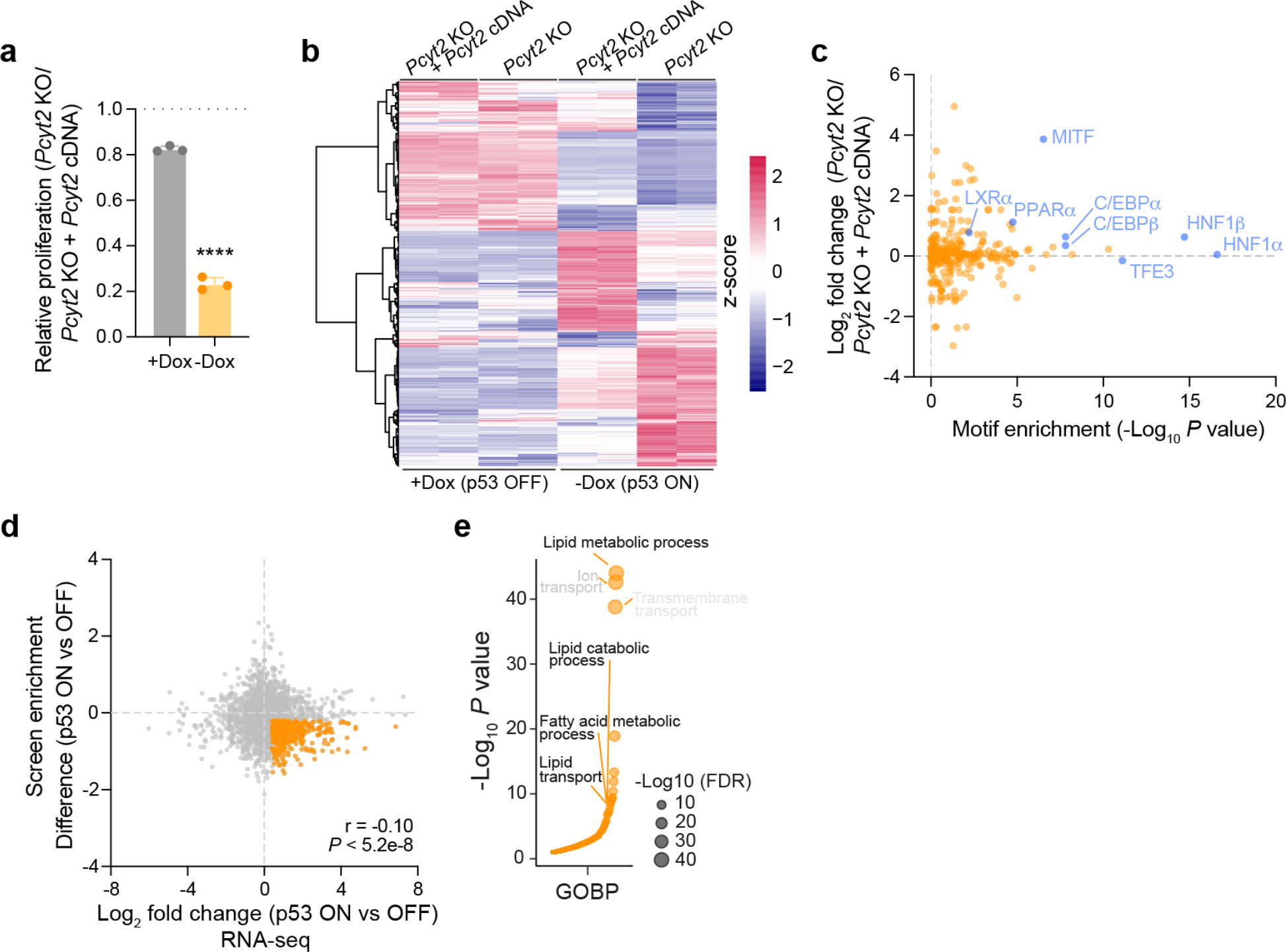
p53 activation increases reliance on genes involved in lipid metabolism. a, Relative proliferation of *Pcyt2* KO cells compared to *Pcyt2* KO + *Pcyt2* cDNA cells cultured on or off dox for 6 days. **b,** Heatmap of differentially expressed genes from RNA-Seq. Included genes exhibit an average FPKM greater than 1 across all conditions and *P* < 0.05 when comparing *Pcyt2* KO to *Pcyt2* KO + *Pcyt2* cDNA cells in the context of p53 activation. See Supplementary Table 1 for gene information. **c,** Scatter plots displaying the significance of transcription factors enriched around the promoters of genes upregulated in *Pcyt2*-knockout compared to *Pcyt2*- knockout + *Pcyt2* cDNA under p53 activation, along with their respective log2 fold changes. **d,** Scatter plot comparing changes in gene expression induced by p53 activation (x-axis, log2 fold change in expression KP^sh^ cells cultured -dox compared to cells cultured +dox) with changes in gene scores following p53 activation (y-axis, gene scores determined by comparing sgRNA abundance of cells following 10 days culture with or without dox). Pearson correlation *r* = -0.10, *P* < 5.2 x 10^-8^. Genes both induced by p53 and more essential following p53 activation (log2 fold change > 0.4; screen enrichment difference < -0.2) are highlighted in orange. Data are available in Supplementary Table 2. **e,** Bubble plot of pathway enrichment for genes highlighted in (**d**).

## References

1. Jackson, B.T. & Finley, L.W.S. Metabolic regulation of the hallmarks of stem cell biology. Cell stem cell 31, 161–180 (2024).

2. Kim, J. & DeBerardinis, R.J. Mechanisms and Implications of Metabolic Heterogeneity in Cancer. Cell metabolism 30, 434–446 (2019).

3. Finley, L.W.S. What is cancer metabolism? Cell (2023).

4. Faubert, B., Solmonson, A. & DeBerardinis, R.J. Metabolic reprogramming and cancer progression. Science 368 (2020).

5. DeBerardinis, R.J. & Chandel, N.S. Fundamentals of cancer metabolism. Sci Adv 2, e1600200 (2016).

6. Luengo, A. et al. Increased demand for NAD(+) relative to ATP drives aerobic glycolysis. Molecular cell 81, 691–707 e696 (2021).

7. Racker, E. Bioenergetics and the problem of tumor growth. Am Sci 60, 56–63 (1972).

8. Bartman, C.R. et al. Slow TCA flux and ATP production in primary solid tumours but not metastases. Nature 614, 349–357 (2023).

9. Warren, L. The biological significance of turnover of the surface membrane of animal cells. Curr Top Dev Biol 4, 197–222 (1969).

10. Kruiswijk, F., Labuschagne, C.F. & Vousden, K.H. p53 in survival, death and metabolic health: a lifeguard with a licence to kill. Nature reviews. Molecular cell biology 16, 393–405 (2015).

11. Boutelle, A.M. & Attardi, L.D. p53 and Tumor Suppression: It Takes a Network. Trends Cell Biol 31, 298–310 (2021).

12. Maddocks, O.D. et al. Serine starvation induces stress and p53-dependent metabolic remodelling in cancer cells. Nature 493, 542–546 (2013).

13. Tajan, M. et al. A Role for p53 in the Adaptation to Glutamine Starvation through the Expression of SLC1A3. Cell metabolism 28, 721–736 e726 (2018).

14. Kastenhuber, E.R. & Lowe, S.W. Putting p53 in Context. Cell 170, 1062–1078 (2017).

15. Morris, J.P. et al. α-Ketoglutarate links p53 to cell fate during tumour suppression. Nature 573, 595–599 (2019).

16. Aird, K.M. et al. Suppression of nucleotide metabolism underlies the establishment and maintenance of oncogene-induced senescence. Cell Rep 3, 1252–1265 (2013).

17. Vassilev, L.T. et al. In vivo activation of the p53 pathway by small-molecule antagonists of MDM2. Science 303, 844–848 (2004).

18. Wishart, D.S. et al. HMDB: the Human Metabolome Database. Nucleic Acids Res 35, D521–526 (2007).

19. Son, Y., Kenny, T.C., Khan, A., Birsoy, K. & Hite, R.K. Structural basis of lipid head group entry to the Kennedy pathway by FLVCR1. Nature 629, 710–716 (2024).

20. Bleijerveld, O.B., Brouwers, J., Vaandrager, A.B., Helms, J.B. & Houweling, M. The CDP-ethanolamine pathway and phosphatidylserine decarboxylation generate different phosphatidylethanolamine molecular species. The Journal of biological chemistry 282, 28362–28372 (2007).

21. Saito, K., Nishijima, M. & Kuge, O. Genetic evidence that phosphatidylserine synthase II catalyzes the conversion of phosphatidylethanolamine to phosphatidylserine in Chinese hamster ovary cells. The Journal of biological chemistry 273, 17199–17205 (1998).

22. Brady, C.A. et al. Distinct p53 transcriptional programs dictate acute DNA-damage responses and tumor suppression. Cell 145, 571–583 (2011).

23. Kenzelmann Broz, D., et al. Global genomic profiling reveals an extensive p53-regulated autophagy program contributing to key p53 responses. Genes & development 27, 1016–1031 (2013).

24. Yamamoto, H., Zhang, S. & Mizushima, N. Autophagy genes in biology and disease. Nat Rev Genet 24, 382–400 (2023).

25. Mizushima, N. & Komatsu, M. Autophagy: renovation of cells and tissues. Cell 147, 728–741 (2011).

26. White, E. Autophagy and p53. Cold Spring Harb Perspect Med 6, a026120 (2016).

27. Wiley, C.D. & Campisi, J. The metabolic roots of senescence: mechanisms and opportunities for intervention. Nat Metab 3, 1290–1301 (2021).

28. Deisenroth, C., Itahana, Y., Tollini, L., Jin, A. & Zhang, Y. p53-Inducible DHRS3 is an endoplasmic reticulum protein associated with lipid droplet accumulation. The Journal of biological chemistry 286, 28343–28356 (2011).

29. Anastasia, I. et al. Mitochondria-rough-ER contacts in the liver regulate systemic lipid homeostasis. Cell Rep 34, 108873 (2021).

30. Vance, J.E. Phospholipid synthesis and transport in mammalian cells. Traffic 16, 1–18 (2015).

31. Vance, J.E. Phospholipid synthesis in a membrane fraction associated with mitochondria. The Journal of biological chemistry 265, 7248–7256 (1990).

32. Zhu, X.G. et al. Functional Genomics In Vivo Reveal Metabolic Dependencies of Pancreatic Cancer Cells. Cell metabolism 33, 211–221.e216 (2021).

33. Lee, B.Y. et al. Senescence-associated beta-galactosidase is lysosomal beta-galactosidase. Aging Cell 5, 187–195 (2006).

34. Tighanimine, K. et al. A homoeostatic switch causing glycerol-3-phosphate and phosphoethanolamine accumulation triggers senescence by rewiring lipid metabolism. Nat Metab 6, 323–342 (2024).

35. Robbins, E., Levine, E.M. & Eagle, H. Morphologic changes accompanying senescence of cultured human diploid cells. The Journal of experimental medicine 131, 1211–1222 (1970).

36. Kurz, D.J., Decary, S., Hong, Y. & Erusalimsky, J.D. Senescence-associated (beta)- galactosidase reflects an increase in lysosomal mass during replicative ageing of human endothelial cells. J Cell Sci 113 **( Pt** **20**), 3613–3622 (2000).

37. Martini, H. & Passos, J.F. Cellular senescence: all roads lead to mitochondria. The FEBS journal 290, 1186–1202 (2023).

38. Serrano, M., Lin, A.W., McCurrach, M.E., Beach, D. & Lowe, S.W. Oncogenic ras provokes premature cell senescence associated with accumulation of p53 and p16INK4a. Cell 88, 593–602 (1997).

39. Dobin, A. et al. STAR: ultrafast universal RNA-seq aligner. Bioinformatics 29, 15–21 (2013).

40. Heinz, S. et al. Simple combinations of lineage-determining transcription factors prime cis-regulatory elements required for macrophage and B cell identities. Molecular cell 38, 576–589 (2010).

41. Love, M.I., Huber, W. & Anders, S. Moderated estimation of fold change and dispersion for RNA-seq data with DESeq2. Genome biology 15, 550 (2014).

42. Lee, T.I., Johnstone, S.E. & Young, R.A. Chromatin immunoprecipitation and microarray-based analysis of protein location. Nature protocols 1, 729–748 (2006).

43. Li, H. et al. The Sequence Alignment/Map format and SAMtools. Bioinformatics 25, 2078–2079 (2009).

44. Zhang, Y. et al. Model-based analysis of ChIP-Seq (MACS). Genome biology 9, R137 (2008).

45. Quinlan, A.R. & Hall, I.M. BEDTools: a flexible suite of utilities for comparing genomic features. Bioinformatics 26, 841–842 (2010).

46. Ramirez, F., Dundar, F., Diehl, S., Gruning, B.A. & Manke, T. deepTools: a flexible platform for exploring deep-sequencing data. Nucleic Acids Res 42, W187–191 (2014).

47. Robinson, J.T. et al. Integrative genomics viewer. Nat Biotechnol 29, 24–26 (2011).

48. Millard, P., Letisse, F., Sokol, S. & Portais, J.C. IsoCor: correcting MS data in isotope labeling experiments. Bioinformatics 28, 1294–1296 (2012).

49. Kumar, A., Mitchener, J., King, Z.A. & Metallo, C.M. Escher-Trace: a web application for pathway-based visualization of stable isotope tracing data. BMC Bioinformatics 21, 297 (2020).

50. Liao, Y., Smyth, G.K. & Shi, W. The R package Rsubread is easier, faster, cheaper and better for alignment and quantification of RNA sequencing reads. Nucleic Acids Res 47, e47 (2019).

